# Low precipitation due to climate change consistently reduces multifunctionality of urban grasslands in mesocosms

**DOI:** 10.1101/2022.09.10.507400

**Authors:** Sandra Rojas-Botero, Leonardo H. Teixeira, Johannes Kollmann

## Abstract

Urban grasslands are crucial for biodiversity and ecosystem services in cities, while little is known about their multifunctionality under climate change. Thus, we investigated the effects of simulated climate change, i.e., increased [CO_2_] and temperature, and reduced precipitation, on individual functions and overall multifunctionality in mesocosm grasslands sown with forbs and grasses in four different proportions aiming at mimicking road verge grassland patches. Climate change scenarios RCP 2.6 (control) and RCP 8.5 (worst-case) were simulated in *walk-in* climate chambers (ca. 7.7 m^2^), and watering was manipulated for normal vs. reduced precipitation. We measured eight indicator variables of ecosystem functions based on below- and above-ground characteristics. Recently established grassland communities responded to higher [CO_2_] and warmer conditions with increased vegetation cover, height, flower production, and soil respiration. Lower precipitation affected carbon cycling in the ecosystem by reducing biomass production and soil respiration. In turn, the water regulation capacity of the grasslands depended on precipitation interacting with climate change scenario, given the enhanced water efficiency resulting from increased [CO_2_] under RCP 8.5. Multifunctionality was negatively affected by reduced precipitation, especially under RCP 2.6. Trade-offs arose among single functions that performed best in either grass- or forb-dominated grasslands. Grasslands with an even ratio of plant functional types coped better with climate change and thus are good options for increasing the benefits of urban green infrastructure. Overall, we provide experimental evidence on the effects of climate change on the functionality of urban ecosystems. Designing the composition of urban grasslands based on ecological theory may increase their resilience to global change.

## Introduction

Urban green infrastructure supports biodiversity and ecosystem services since it serves as habitat, regulates temperature and stormwater, improves air and water quality, and increases aesthetics [1]. With increased anthropogenic [CO_2_], higher temperatures, changed precipitation, and more extreme events [2], functional urban ecosystems are relevant for mitigating climate change via ecosystem services [3–5], while also threatened by climatic extremes [6,7]. The various components of climate change can alter the physiological response of organisms, the dynamics of populations, and the development of communities, ultimately affecting ecosystems’ functions and services [8]. In grasslands, for example, higher temperature increases ecosystem respiration and reduces productivity [9], while elevated [CO_2_] increases soil respiration [10] and above-ground productivity [11]. In turn, water stress affects net ecosystem CO_2_ exchange [12,13], and increases soil water retention, nutrient leaching, and soil fertility [12].

Grasslands represent a high proportion of urban ecosystems [14]. Among those, urban lawns are the most frequent components of green infrastructure [15]. However, due to irrigation, regular mowing, and fertilization, urban lawns have become less diverse [16], and have altered biogeochemical cycles [17,18]. With growing initiatives to replace intensively managed lawns in cities with diverse meadows [19,20], understanding their functioning and resilience to climate change will allow for better design and more effective restoration measures of green urban spaces. If adequately implemented, urban grasslands can harbor large numbers of plant species [21], offer habitat and food for animal species [22,23], and provide numerous ecosystem services to humans [1].

A deeper understanding of the biotic drivers of the multiple functions performed by urban grasslands and their responses to climate change helps identify synergies and trade-offs among ecosystem functions, with contrasting consequences to ecosystem services [24]. During the past decades, various studies highlighted the role of taxonomic and functional diversity on single ecosystem functions and ecosystem multifunctionality [24,25], particularly in experimental or semi-natural grasslands [26]. Besides species richness, functional diversity, and the composition of different functional types can explain ecosystem functioning [27,28]. In grasslands, grasses and forbs are useful surrogates of functional diversity, depicting specific adaptations to the physical environment, patterns of resource use, and resilience to disturbances [27,29], also under climate change [30]. Therefore, variation in the abundance of grasses and forbs might translate into changes in grassland functioning.

Despite increasing evidence that community composition affects the resilience of grassland functioning to climate change, most studies focus only on species richness as a driver of resilience to climate change [but see 31]. Moreover, few studies consider the combination of different climate change components [e.g., 32,33]. Thus, the effects of climate change on varying compositions of grasslands remain poorly understood [34], and hamper the recognition of relevant aspects to the design and management of multifunctional urban grasslands.

Here, we evaluate whether the simultaneous impacts of simulated climate change and functional type composition affect the multifunctionality of urban grasslands using mesocosms in climate chambers as model systems. To address current research gaps, we assess the combined effects of three components of climate change, i.e., higher [CO_2_], increased temperatures, and reduced precipitation on the multifunctionality of experimental grasslands composed of contrasting proportions of forbs vs. grasses. We expected: (i) higher temperature and [CO_2_] to enhance carbon cycling of mesocosm grasslands; (ii) reduced precipitation to decrease vegetation performance and ecosystem multifunctionality; (iii) diverse functional type composition to produce contrasting effects on grassland functioning under current environmental conditions, with climate change enhancing specific responses of grass- and forb-dominated communities; and (iv) mesocosm grasslands with high evenness of plant functional types to exhibit higher multifunctionality under climate change.

## Materials and methods

We implemented a mesocosm experiment imitating early phases of grassland establishment in urban settings, assuming the sowing of target species in prepared bare soil of urban road verges (S1 File). In this experiment, we manipulated the relative abundance of forbs and grasses to assess grassland functioning in response to variation in plant functional types and simulated climate change (decreased precipitation, increased temperature, and [CO_2_]). We measured different indicator variables to assess the functioning of grasslands: vegetation structure and productivity, soil, and water regulation as proxies of functions related to ecosystem services expected from multifunctional urban ecosystems.

### Design of experimental grassland communities

We tested four grassland mixtures: A seed mixture of only grasses, representative of species-poor lawns. Thereupon, the communities were complemented by a mixture of native forbs used in restored urban road verges [35] to reach the following proportions of forbs and grasses, respectively, i.e., 0:100 (F0), 50:50 (F50), 75:25 (F75), and 100:0 (F100). We selected five native grass and 26 forb species (five being legumes) occurring in urban areas of Bavaria, following criteria relevant to urban pollinators and plant performance in urban contexts (S1 Table). Mixture F0 comprised five grass species (one plant family), F100 contained 26 species (12 families), and F50 and F75 had 31 species (13 families). Seed mixtures were assembled by weight to a desired density of 4400 seeds m^-2^. The contribution of each species to the seed mixture was adjusted to each treatment, whereby each species of the respective functional type had the same proportion (S2 Table). The actual establishment of the experimental communities was not recorded, but it looked similar to the respective mixtures. Because of focusing on the influence of the plant functional types (grasses vs. forbs), species numbers were not accounted for, but considered minor when assessing the effects of functional composition on the grasslands.

### Experimental design and monitoring

To test the effects of plant functional types proportions (‘forb proportion’ henceforth) and climate change on urban grassland multifunctionality, we used a 10-week experiment in four walk-in climate chambers (2.4 × 3.2 × 2.2 m^3^; area 7.7 m^2^; S1 Fig) located at the TUMmesa facility of Technical University of Munich (Freising, Germany). We simulated environmental conditions of two IPCC climate change scenarios (RCP 2.6 and RCP 8.5; two chambers per scenario) reflecting environmental parameters of May–July when vegetation development is fast. For RCP 2.6 (control), with temperature and [CO_2_] similar to current values, we reproduced median hourly values of temperature between 2000 and 2019 [36] and [CO_2_] recorded in central areas of Munich with a TDLAS system [37]. For RCP 8.5 (worst-case scenario), median values of the record warm years for C Europe 2003, 2015, and 2018 were used, while the values of [CO_2_] were set to double the current ones. Simulation of the worst-case climate change scenario (RCP 8.5) increased [CO_2_] and air temperature by 486 ppm (+99.7%) and 3.1 °C (+17.6%), respectively. The relative humidity was lower under RCP 8.5 (–4.7%). Thus, the realized simulation of temperature and [CO_2_] lies within the expectations under RCP 8.5 by the end of the century [2], and allowed for clear differentiation between the two scenarios manipulated in the climate chambers (S2 Fig).

Sixty-four plastic mesocosms (70 × 40 × 23 cm^3^ (L x W x H)) were filled with a mixture of washed sand:potting substrate (70:30) in the lowest 10 cm, and urban lawn substrate in the upper 10 cm, to mimic the soil depth conditions of urban road verges (Fig 1). Each mesocosm had six 1-cm diameter holes in the bottom to allow water drainage, and a plastic container placed below to collect the drained water. Mesocosms were distributed on floodable tables (four per table). Four tables were allocated within each climate chamber. A grassland mixture (F0, F50, F75, or F100) was randomly assigned to each mesocosm on a table, while the two simulated precipitation regimes were randomized among the tables in each chamber. Thus, each mesocosm represented a combination of seed mixture, watering regime, and climate change scenario ([CO_2_] and temperature) (S3 Fig).

**Fig 1.**
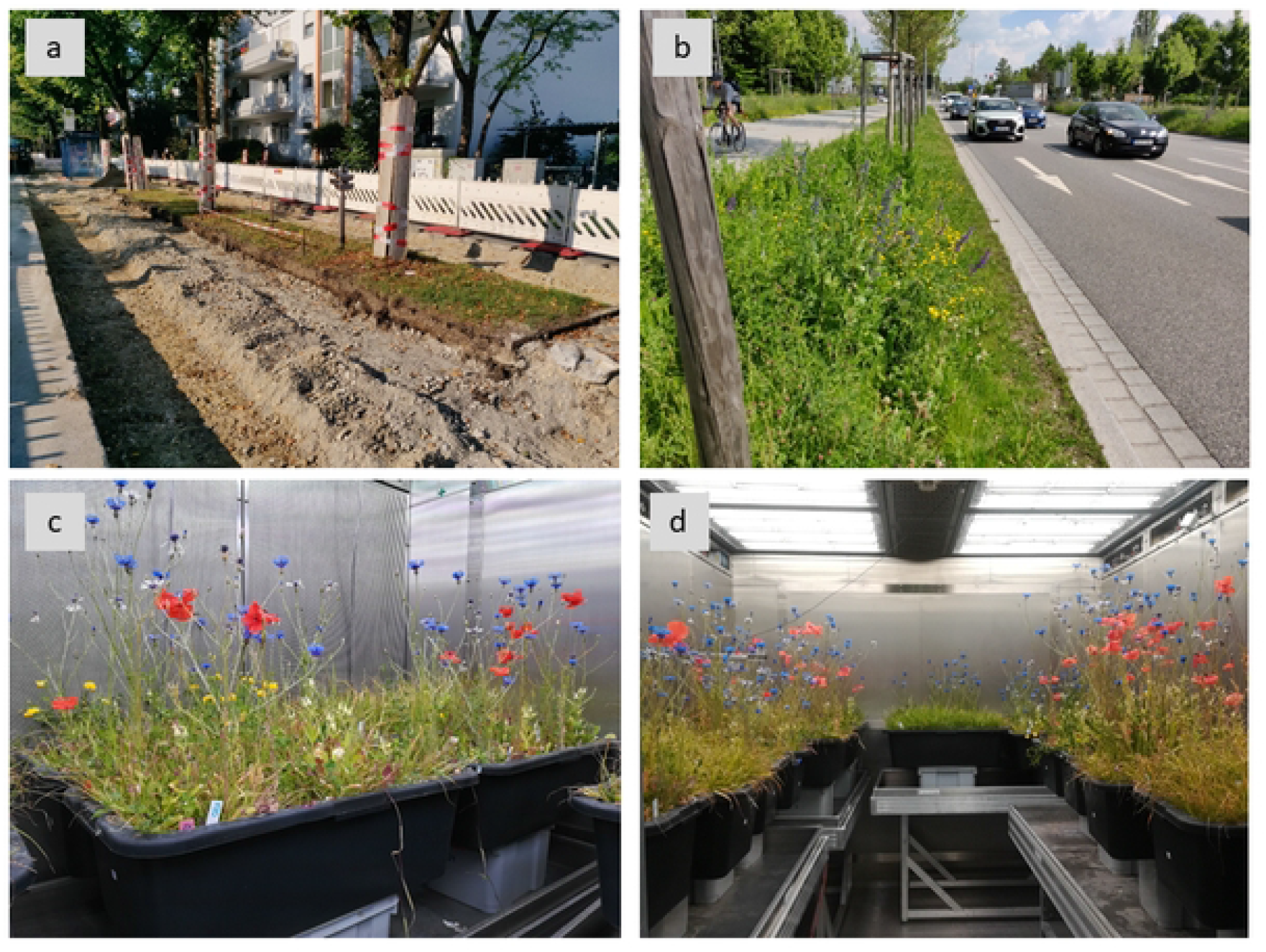
Urban road verge grasslands mimicked in mesocosms in climate chamber experiment. (a) Road grasslands often occur on heavily sealed ground with shallow soil and using few grass species. (b) Restored grasslands are species-rich and adapted to urban conditions to increase ecosystem service delivery. (c) In the mesocosm experiment, species used for restoring urban road verges were sown, and the communities developed under similar conditions (e.g., substrate characteristics, limited soil depth, and surrounding space). (d) Mesocosm grasslands of varying functional compositions were submitted to climate change scenarios. Pictures (a) and (b) taken in Munich, Germany (courtesy of Simon Dietzel); pictures (c) and (d) were taken at weeks 9 and 10 of mesocosm development.

Each seed mixture was thoroughly spread by hand onto the prepared substrate of the mesocosms, which were kept well watered during the first 27 days after sowing. Precipitation was simulated by watering the mesocosms from above every other day with a hose sprayer. We adopted the mean precipitation recorded in Munich from May to July between 2000 and 2019. For the reduced precipitation treatment, we halved the amount of water in the mesocosms, matching the mean precipitation amounts recorded in the extremely dry years 2003, 2015, and 2018. There was no fertilization during the experiment.

We measured indicator variables of ecosystem functions of interest for urban greening at each mesocosm. The measurements represented above- and below-ground compartments of urban grasslands, with which we approximate ecosystem functions [26,38–40]. A comprehensive overview of the indicator variables, experimental measurement, their role in urban ecosystem services, and potential responses to the manipulated components of climate change is available in S3 Table. We recorded five variables related to vegetation (i.e., above- and below-ground biomass, flower production, plant cover, and plant height), constituting a good proxy for productivity and determining to a significant extent ecosystem functioning [41]. Moreover, we considered cover and height as structural components of vegetation that, besides productivity [41], relate to above-ground space filling, habitat offer, and soil protection [26,41]. We also measured soil respiration as it correlates to C exchange and represents the largest component of ecosystem respiration [42]. Finally, water retention and water loss through evapotranspiration (ET) were measured using a watering and weighing protocol adopted from studies on green roofs [43,44], and used as a proxy for rainwater regulation functions of urban grasslands in shallow road verges. For ideal stormwater regulation and transpirational cooling support, urban grasslands need to use high amounts of water when available after a rain event, and persist during periods of water scarcity [43,45].

### Data analyses

Before assessing single indicator variables, above- and below-ground biomass and floral abundance were calculated per area. We averaged measures taken multiple times in each mesocosm (e.g., soil respiration), so every variable was included only once in the analyses. Water retention and loss were expressed as retained or lost water fractions (values between 0 and 1).

We calculated two indices of ecosystem multifunctionality with the eight indicator measurements: the ‘averaging’ and ‘single threshold approaches’, using the protocol of [25] and [24], and implemented both with the R package ‘Multifunc’ version 0.8.0 [25]. Data exploration for collinearity resulted in a strong correlation between above-ground biomass and floral density (above |0.7|; S4 Fig; [46]). Therefore, we calculated each index twice, once without above-ground biomass and another without floral density. Because floral density and above-ground biomass are essential indicators of the expected functioning of urban grasslands, we calculated two values per multifunctionality approach instead of discarding one of these indicator variables. Hierarchical clustering following [24] was used before calculating the two approaches to multifunctionality to balance calculations among correlated functions by identifying subsets of related variables (S5 Fig). This procedure prevented closed-related variables from disproportionally driving multifunctionality to certain aspects of ecosystem function [24]. After clustering of the variables, an optimal number of clusters was determined following the ‘elbow method’, and the variables within each cluster were weighed equally by giving a proportional weight summing up to one for each cluster. Three clusters were determined and the weighted indicators variables contributed to each multifunctionality calculation.

In the averaging approach, all standardized values of the weighed indicator variables were averaged per mesocosm, and this measure of multifunctionality was modeled later against climate change components and forb proportion. In the threshold approach, we selected a 70% threshold as a desired level of multifunctionality for urban road verge grasslands with high responsiveness [47]. To obtain the maximum observed value of each assessed indicator variable [25], we averaged the six highest observations per indicator variable (top 10% values) to reduce outlier effects [25,48], and to reflect conservative maximum multifunctionality attained by the urban grassland systems we studied in the mesocosms. In the calculation, each ecosystem function exceeding 70% of the standardized maximum contributed to the multifunctionality score with its respective weighted value obtained after clustering. In light of the challenges imposed by climate change to the establishment of functional urban ecosystems, we considered this threshold to be a reasonable assumption desired for these systems. Finally, because of each approach’s known advantages and disadvantages in measuring multifunctionality [25], we presented and discuss the results based on the assessment of individual measurements.

We tested the effects of forb proportion, precipitation, and RCP scenario on single indicator variables and the two metrics of multifunctionality calculated for the experimental grasslands. We considered full models with all possible two-way interactions and conducted model simplification until at least the main fixed effects remained in the model. All possible models were compared via the dredge function [‘MuMIn’; [49]], and we selected the simplest ones among those with a difference in AICc < 2. For continuous data (i.e., above-ground biomass, below-ground biomass, plant cover, plant height, respiration, and the multifunctionality indices), linear mixed-effects models were used [‘lme4’; [50]]. Data stemming from the measurement of water regulation (i.e., water retention and loss) were analyzed using beta regression [‘glmmTMB’; [51]]. For the floral density, we fitted a zero-inflated negative binomial mixed-effects model (ZIP) after testing and detecting an excess of zeros in the data using a simulation approach and comparing the proportion of zeros in the data to the simulated ones expected based on a Poisson model [52]. We used generalized linear mixed-effects models because they provide a flexible approach to model non-normally distributed data (i.e., counts, fractions) and the crossed character of the data in the experimental design. In all cases, we included the identity of climate chambers as a random effect. The fit of the models was checked with the DHARMa package [52]. Post-hoc analyses were applied to compare functioning between levels of forb proportion whenever a significant effect was found and to correct for multiple comparisons [53]. Data analysis was conducted with R version 4.1.2 [54].

## Results

### Response of single functions

The proportion of forbs and the simulated precipitation additively influenced above-ground biomass (Table 1; Fig 1a). Communities with forbs produced more aboveground biomass across environmental conditions than grass-only communities (F100, t = 5.02, p < 0.001; F75, t = 3.49, p < 0.001; F50, t = 4.09, p < 0.001; complete model outputs in S4 Table). Reduced precipitation negatively influenced above-ground biomass production in all communities (t = - 3.83, p < 0.001; S4 Table), regardless of climate change scenario. Even though above-ground biomass was higher under warmer temperatures and higher [CO_2_], no statistical evidence was observed for an RCP scenario effect (Table 1; S4 Table). Below-ground biomass was additively affected by forb proportion and precipitation (Table 1, Fig. 1b). Opposite to above-ground biomass, grass-only communities (F0) presented highest root biomass under all climatic conditions (S4 Table). Communities with even functional composition (F50) had higher belowground biomass than forb-only communities (t = 2.93, p = 0.025; S5 Table). Reduced precipitation negatively affected root biomass production in all experimental communities (t = -1.96, p = 0.055).

**Table 1.** **Effects of forb proportion, precipitation, RCP climate change scenarios, and their interactions on eight indicator variables and multifunctionality of mesocosm grasslands in climate chambers**.

| Indicator variable | Forb proportion |  | RCP scenario |  | Precipitation |  | RCPscen:Precip |  | RCPscen:Forb prop |  | Precip:Forb Prop |  |
| --- | --- | --- | --- | --- | --- | --- | --- | --- | --- | --- | --- | --- |
|  | Statistic | p-value | Statistic | p-value | Statistic | p-value | Statistic | p-value | Statistic | p-value | Statistic | p-value |
| Above-ground biomass | 28.82 | **<0.001** | 1.49 | 0.22 | 14.64 | **<0.001** |  |  |  |  |  |  |
| Below-ground biomass | 58.76 | **<0.001** | 0.99 | 0.32 | 3.85 | **0.05** |  |  |  |  |  |  |
| Plant cover | 12.87 | **0.005** | 13.90 | **<0.001** | 47.84 | **<0.001** |  |  |  |  |  |  |
| Plant height | 6.38 | 0.09 | 6.68 | **0.010** | 1.45 | 0.23 | 60.56 | **<0.001** | 27.04 | **<0.001** | 16.66 | **0.001** |
| Floral density | 179.82 | **<0.001** | 6.69 | **0.010** | 17.99 | **<0.001** |  |  |  |  |  |  |
| Soil respiration | 4.63 | 0.20 | 61.81 | **<0.001** | 18.83 | **<0.001** |  |  |  |  |  |  |
| Water retention | 0.44 | 0.93 | 0.57 | 0.45 | 29.02 | **<0.001** | 8.02 | **0.005** |  |  |  |  |
| Water loss (ET) | 4.09 | 0.25 | 1.72 | 0.19 | 26.28 | **<0.001** | 6.84 | **0.009** |  |  |  |  |
| Avg. Multifunctionality-AGB | 2.07 | 0.56 | 0.08 | 0.78 | 9.44 | **0.002** | 4.25 | **0.039** | 12.79 | **0.005** |  |  |
| Avg. Multifunctionality-Flowers | 1.05 | 0.79 | 0.11 | 0.74 | 10.22 | **0.001** | 5.35 | **0.021** | 11.52 | **0.009** |  |  |
| 70% Multifunctionality AGB | 2.19 | 0.53 | 6.77 | **0.009** | 14.75 | **<0.001** |  |  |  |  |  |  |
| 70% Multifunctionality-Flowers | 3.15 | 0.37 | 12.17 | **<0.001** | 14.82 | **<0.001** | 9.92 | **0.002** |  |  |  |  |

Responses were analyzed with (generalized) linear mixed-effects models. Climate chambers were set as random factors. Shown are
outputs of ANOVA tables displaying the significance of the tested drivers after model simplification. Degrees of freedom are forb
proportion (df = 3), precipitation (df = 1), RCP scenario (df = 1), RCP scenario x precipitation, forb proportion x RCP scenario (df =
3), and forb proportion x precipitation (df = 3). Significance set at $p < 0.05$ depicted in bold (n = 4).

Floral density significantly responded to forb proportion, precipitation, and RCP scenario (Table 1, Fig 1c). Density of flowers increased with forb proportion (t = [10.11, 12.54], p < 0.001), more flowers were produced under normal precipitation (t = 4.24, p < 0.001), and scenario RCP 8.5 (t = 2.59, p = 0.009). Plant cover was additively influenced by forb proportion, precipitation, and climate change scenario (Table 1, Fig 1d). Plant cover increased with lower forb proportion (t = [2.07, 2.55], p < 0.05; S4 Table), and under RCP 8.5 (t = 3.73, p = 0.06), but decreased with reduced precipitation (t = -6.92, p < 0.001). Forb proportion, precipitation, climate change scenario, and all two-way interactions affected mean plant height (Table 1, S4 Table, Fig 1e). All communities containing forbs were taller than grass-only communities under normal precipitation conditions, while there were no clear differences under reduced precipitation (S6 Table). Plant height was negatively affected by the interaction of reduced precipitation and scenario RCP 8.5 (t = 7.78, p < 0.001), most likely due to insufficient water supply for stimulating growth, even though higher [CO_2_] increases the fixation of C and plant growth. Under RCP 8.5, grass-only communities were shorter than all other functional compositions (S6 Table), and grasslands with an even composition of forbs and grasses (F50) were significantly taller than forb-dominated grasslands (i.e., F75 and F100). Climate change scenario and precipitation additively affected soil respiration (Table 1). Soil respiration rate was lower in mesocosm grasslands with reduced precipitation (t = -4.34, p < 0.001), and under scenario RCP 8.5, soil respiration increased (t = 7.86, p = 0.016; Fig 2e). Water regulation did not respond to forb proportion and RCP scenario (Table 1). In turn, both indicator variables of water regulation were affected by the interaction between precipitation and RCP scenario (Table 1). Under RCP 2.6, water retention substantially increased in grasslands with reduced precipitation, whereas water loss decreased significantly when precipitation was reduced (Fig 3b; S4 Table).

**Fig 2.**
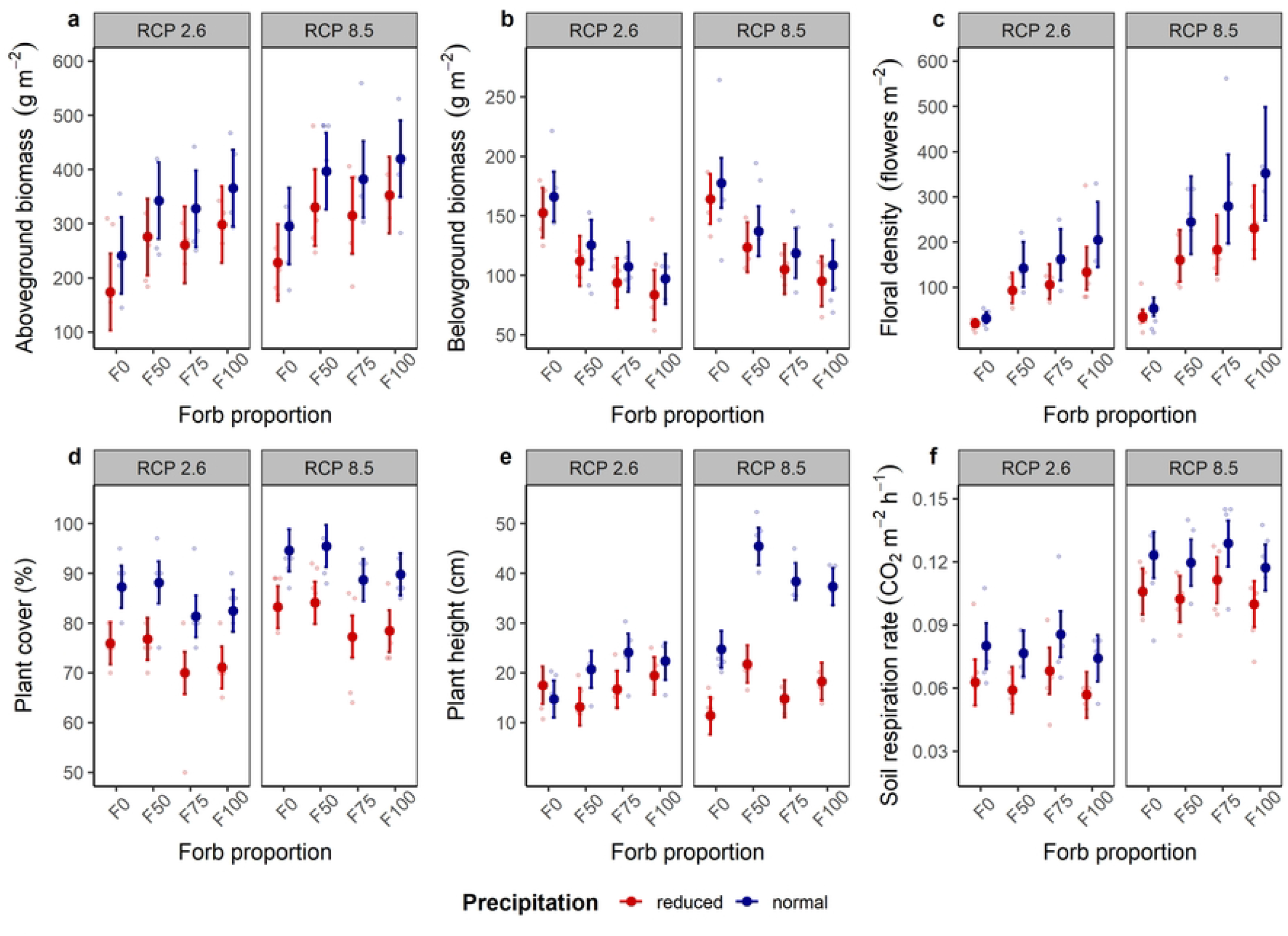
Effects of forb proportion, precipitation, and climate change scenarios on indicators of grassland functioning. (a) Aboveground biomass, (b) belowground biomass, (c) floral abundance, (d) vegetation height, (e) cover, and (f) soil respiration rate (means ± 95% confidence intervals) of mesocosm communities in climate chambers. Data points are light-colored circles (n = 4).

**Fig 3.**
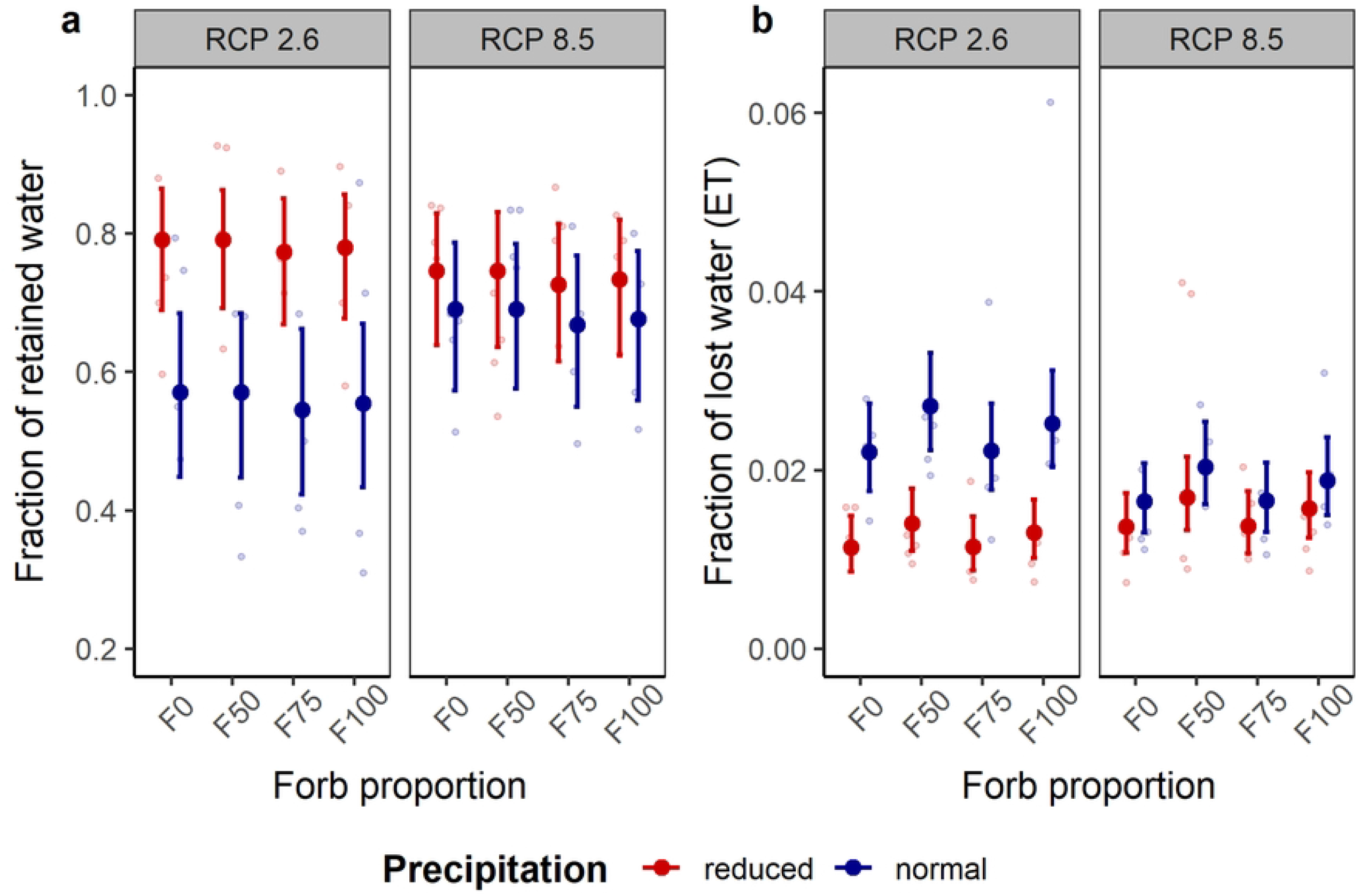
Effects of forb proportion, precipitation and climate change scenarios on water regulation in grasslands. (a) Fraction of retained, and (b) lost water by evapotranspiration [ET] (means ± 95% confidence intervals) of mesocosm communities in climate chambers. A heavy-rain event was simulated to derive information on water regulation function. Data points are light-colored circles (n = 4).

### Urban grassland multifunctionality

Precipitation and its interaction with RCP scenario affected averaged multifunctionality (Table 1). Decreased precipitation was overall detrimental for averaged multifunctionality, whereas RCP 8.5 in interaction with normal precipitation was beneficial for averaged grassland multifunctionality (S7 Table, Fig 4a,c). Under RCP 8.5, grasslands evenly composed of grasses and forbs (i.e., F50) presented higher averaged multifunctionality than grass-only communities in both cases, i.e., considering the calculations using above-ground biomass and floral density (t = 3.40 and 3.29, p = 0.001 and 0.002, respectively; S7 Table). F75, in interaction with RCP 8.5, positively affected multifunctionality when floral abundance was used instead of above-ground biomass for multifunctionality calculation (t = 2.05, p = 0.045), underscoring the biotic control of multifunctionality under climate change.

**Fig 4.**
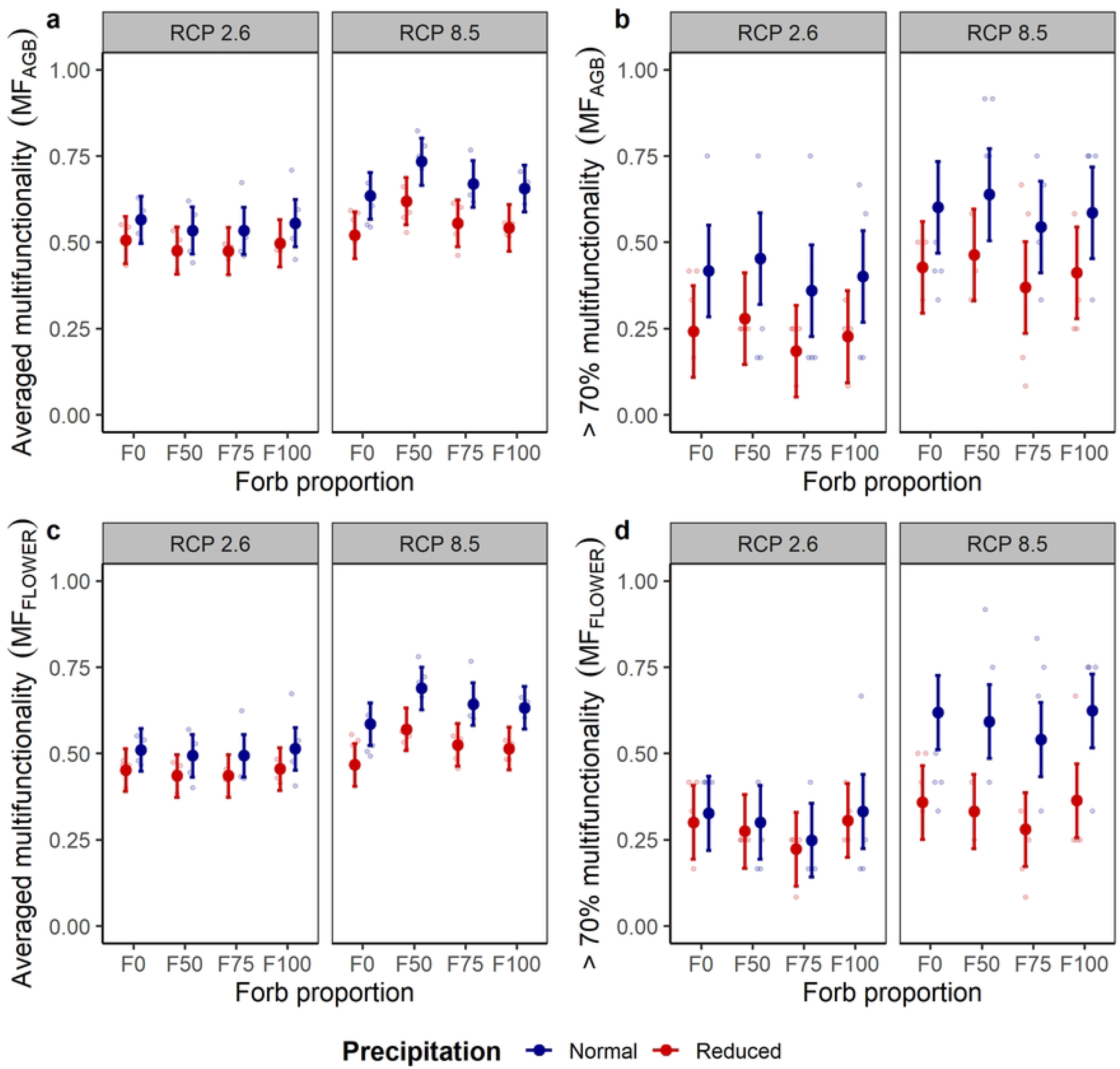
Effects of forb proportion, precipitation and climate change scenario on grassland multifunctionality. Two approaches to multifunctionality (MF) were calculated using eight indicator variables, including either above-ground biomass (a, b) or floral density (c, d) of mesocosm communities in climate chambers. Averaged standardized values of the individual indicators contributed to averaged multifunctionality, whereas for the selected threshold, each function indicator exceeding 70% of the standardized maximum contributed to multifunctionality. Shown are means ± 95% confidence intervals. Data points are light-colored circles (n = 4).

Only precipitation affected the functioning of mesocosm grasslands at or above a threshold of 70% multifunctionality (Table 1). We observed more ecosystem functions above this threshold under normal precipitation (Fig. 4b,d, S7 Table). Moreover, when multifunctionality at 70% threshold considered flower density instead of above-ground biomass, the interaction between precipitation and RCP scenario became significant, with a negative effect of RCP 8.5 interacting with reduced precipitation (S7 Table; Fig 4d). At lower levels of multifunctionality (e.g., 50% threshold), decreases in precipitation were consistently detrimental. At the same time, functional composition in interaction with RCP scenario remained important for increased multifunctionality, particularly in communities with even compositions of forbs and grasses, i.e., F50 (S2 File).

## Discussion

Understanding the functioning of urban grassland is critical for climate change adaptation, the delivery of ecosystem services, and biodiversity protection in cities. Here, we assessed ecosystem functioning under climate change in mesocosms, mimicking early establishment of road verge grasslands. Our assessment of single indicator variables and multifunctionality emphasizes the importance of designing urban grasslands informed by ecological theory in times of climate change.

### High [CO_2_] and temperature enhance carbon cycling during grassland establishment

RCP scenario (i.e., air temperature and [CO_2_]) affected carbon-cycling processes and, thus, crucial functions for multifunctional urban grasslands. These results confirm the conclusions of other climate change experiments [55]. RCP 8.5 positively affected individual functions like vegetation cover, mean height, floral abundance, and soil respiration. Therefore, attributes related to vegetation growth and habitat availability for higher trophic levels benefit from warmer and CO_2_-richer conditions. In addition, increased soil respiration under RCP 8.5 indicates an increased metabolism in the soil accelerating C turnover driven by increased plant photosynthesis, and a higher release of recently allocated C via respiration [42,55], especially under non-water limiting conditions.

Despite RCP 8.5 favored water-use efficiency and indicator variables of productivity were greater even in water-limited mesocosm grasslands (i.e., under reduced precipitation), the interaction between RCP 8.5 and normal precipitation was positively associated with higher grassland multifunctionality. For single functions and multifunctionality, elevated temperature and CO_2_ increased multifunctionality if water supply suffices for the greater water amount required to sustain higher plant growth. This indicates warmer and CO_2_-richer conditions favored plant development [56] due to CO_2_ fertilization [57], particularly in the early stages of community assembly [31].

Since photosynthesis and respiration represent the two paramount fluxes of CO_2_ in ecosystems linked to C allocation, increased cycling of C can be assumed from climate-change conditions such as those in RCP 8.5, stimulating plant growth and soil respiration [10]. Plants under RCP 8.5 assimilate more C [57] and allocate more of the photosynthesized C below-ground, leading to increases in soil respiration [42], because warmer temperatures accelerate metabolic activity in the soil. While urban lawns have higher respiration rates and contribute to urban CO_2_ emissions [17], the respiration rates recorded in our experiment were lower than reported for intensively managed urban grasslands, even under RCP 8.5 [17]. This observation suggests the relevance of management measures rather than grassland composition and climate change conditions for the disproportionate enhancement of CO_2_ efflux in urban grasslands [58]. Thus, we argue that management-related drivers of exacerbated soil respiration (frequent mowing, fertilization, and irrigation) can be a critical factor for functioning urban grasslands in terms of C exchange and the expected role of urban grasslands in climate change mitigation in urban areas.

### Detrimental role of reduced precipitation on urban grassland functioning

Precipitation was a consistent driver of grassland multifunctionality across the two approaches assessed. As predicted for the growing season in Central Europe [2], lower precipitation negatively affected all functions related to carbon cycling and ecosystem multifunctionality. Less precipitation has more detrimental effects on carbon fixation in plant tissues (shoots, flowers, roots) than in ecosystem respiration, thus reducing net C uptake in ecosystems [42]. Previous studies also reported strong effects of rain reduction on grassland multifunctionality [13], underlining the challenging influence of modified precipitation patterns, especially in the above-ground productivity of ecosystems [55]. In urban areas, the effects of reduced precipitation might be enhanced because soil sealing and building materials intensify water loss and stress [7]. Despite studies reporting the stabilizing effects of increased species richness and adapted composition on grassland responses to drought [13,27,30], our findings did not support biotic control in multifunctionality under reduced precipitation. Moreover, decreased precipitation was detrimental for averaged multifunctionality under RCP 8.5, likely because the water-saving effect of elevated [CO_2_] did not compensate for the temperature-driven drying of the soil that negatively affected water supply for plant metabolism and grassland functioning.

With reduced precipitation, water retention capacity in the grasslands increased, while RCP scenario modulated this effect. Water-stressed grasslands used more water and generated more space in the soil and plant tissues for water retention between rain events, similarly to more engineered urban ecosystems (e.g., green roofs; [44]). Notwithstanding, less water retention occurred under RCP 8.5, given the increased efficiency in water use by plants with elevated [CO_2_], preventing high water absorption by plants, and reducing soil water retention capacity following a rain event. In contrast, reduced precipitation reduced water circulation via ET, which supports transpirational cooling in cities [59,60]. Particularly under RCP 2.6, we found a steeper decrease in water loss via ET, because plants tended to keep absorbed water to sustain their metabolism. Instead, in communities growing under climate change conditions (RCP 8.5), losses in water did not substantially change due to CO_2_ fertilization that increases water use efficiency by plants.

### High evenness in functional type composition is beneficial for multifunctionality but not for all individual functions

The functional composition of mesocosm grasslands (i.e., the proportion of forbs and grasses) was a modulating driver of multifunctionality. Under RCP 8.5, communities with an even ratio of plant functional types (F50) exhibited higher average multifunctionality. They favored some individual indicator variables compared to other compositions (e.g., below-ground biomass and plant height and cover). This finding underscores the biotic control of multifunctionality under climate change, and supports the complementarity between plant functional types as one aspect explaining improved functioning of grasslands [28]. Therefore, differences in composition among grasses and forbs may better explain multifunctionality during the early assembly of urban grasslands, reinforcing the need to account for functional types when assessing ecosystem function responses to climate change [29,30].

Nevertheless, some functions were strongly correlated with the dominance of either forbs or grasses, indicating their prominent role in specific ecosystem processes. Below-ground biomass was higher in grass-only communities and decreased steadily with a lower abundance of grasses in the community. The higher ability of grasses to allocate biomass to the roots and produce an intricate network to capture and accumulate resources, especially in the upper soil layers, confirms [18]. Similarly, grasses also favored vegetation cover, highlighting their ability to cover bare soil rapidly [15] and support surface cooling [59]. In contrast, forbs increased above-ground biomass, vegetation height, and floral abundance, significantly impacting carbon fixation, and habitat and resources for animals, especially arthropods [20]. This contrast in functions represents trade-offs in functioning that might reflect the degree of multifunctionality expected from urban grasslands. Tall-grass meadows might maximize habitat and aesthetics [16,20]; grass-dominated ones might, in turn, optimize functioning as runoff controllers [60], soil stabilizers, and recreation areas [18]. Such trade-offs among functions might explain why our averaged multifunctionality assessment underscores that high evenness between grasses and forbs (F50) increases grassland multifunctionality under climate change.

We did not observe differential effects of functional composition on water regulation. Since urban lawns increase water capture and infiltration because their root system enables extensive soil exploration [60,61] and the opening of channels [62], we anticipated grass-only (F0) communities to retain more water. Despite not being statistically different, communities with even functional composition (F50) retained greater amounts of water. Presumably, a high complementarity and evenness in the communities’ above- and below-ground traits can be beneficial for water capture and infiltration [63]. Furthermore, we observed the lowest water losses in grass-only communities because grasses efficiently withstand water stress [62]. In contrast, communities with even functional composition (F50) lost more water via ET and thereby contributed to water circulation in the ecosystem, one desired function of green urban areas where runoff due to sealing is high [60]. Despite water regulation not being controlled by functional composition in our experiment, evenly composed communities (F50) tended to show a greater capacity to capture water and circulate it via ET in urban grasslands.

Since a large body of literature demonstrates that multifunctionality increases with plant diversity [26,38,48,64], we did not address plant diversity within this study: First, we were not interested in understanding diversity effects explicitly. We sowed five (F0), 26 (F100), or 31 (F50, F75) species in our mesocosm grasslands, exemplifying real-life mixture diversity used in urban grasslands and not representing an experimentally intended gradient of plant species diversity. Second, we did not aim to disentangle species-richness influences from functional-composition ones but to understand grassland functioning in response to climate change as mediated by its functional composition. However, we cannot rule out the confounding effects of species richness and functional composition in our setup. Nevertheless, two of the tested mesocosm grassland compositions that reflected the highest richness of sown species (31 plant species) contained the two functional types of interest and revealed contrasting multifunctionality. The mixture with high evenness between forbs and grasses (F50) performed best for most functions. In contrast, F75, also containing 31 sown species, showed lower levels of (multi)functionality. This suggests that besides diversity and functional type richness, the dominance or presence of specific functional types strongly influences ecosystem multifunctionality [65], and that in addition to plant species richness, the effects of functional composition and individual species on ecosystem functioning, e.g., biomass production [28,64,65] needs to be accounted for.

A strength of our experiment was the highly controlled factors mimicking the limited available soil volume and the strong edge effect experienced by the grasslands of road verges. Even though a complete set of factors related to global change could not be realized, our full-factorial combination of projected changes helps understanding ecosystem responses to climate change [66]. However, we acknowledge that mesocosm experiments represent a trade-off with external validity of responses given missing realistic conditions [66], such as disturbances and interactions experienced in natural systems. Moreover, we cannot extrapolate long-term responses of grassland multifunctionality under climate change. For example, biomass production responses to CO_2_ fertilization may stabilize over time [31,66,67], whereas we explicitly provide evidence of responses of recently established road verge grasslands to climate change.

The importance of the drivers of grassland multifunctionality differed between the two approaches applied in our assessment, likely due to the metric used for its evaluation [25]. The averaging approach depended mainly on the functions with large absolute values responding to factors tested [26]. For instance, ecosystem productivity was controlled by the functional composition of the communities (i.e., forb proportion), and a similar control was found in averaged multifunctionality. In turn, we considered the 70% threshold as a reasonable assumption desired for the multifunctionality of urban grasslands, even though additional challenges and disturbances faced under heavily urbanized settings may imply the consideration of lower thresholds of multifunctionality is necessary. In fact, including different threshold levels when assessing the multifunctionality of urban grasslands might inform management and restoration decisions due to providing an integrative assessment of the functioning of these systems. Overall, with the two approaches to multifunctionality deployed, we found high consistency in the adverse effects of reduced precipitation on the functioning of recently established road verge grasslands. The assessment of multifunctionality in restored urban grasslands and possible derived ecosystem services should be encouraged to increase their ecological value in cities.

## Conclusions

The study provides valuable insights into the functioning of road verge grasslands under climate change conditions. They highlight that reduced precipitation is detrimental for functioning of urban grasslands. At least during the early stages of their development, grasslands can benefit from warmer and CO_2_-richer conditions that foster productivity. Moreover, our averaged multifunctionality assessment underscores the role of functional composition of urban grasslands in modulating the responses to a changing environment. At the same time, the threshold level considered provides a realistic approach to multifunctionality. Urban grasslands with a high evenness between grasses and forbs are suitable for obtaining higher levels of multifunctionality, and are more effective in biodiversity protection and ecosystem services in times of climate change.

## ACKNOWLEDGMENTS

Climate data (multi-annual air temperature in Munich) were provided by Deutscher Wetterdienst (DWD). The Chair of Environmental Sensing and Modeling of the TUM kindly provided data on [CO_2_] recorded in Munich. Roman Maier supported the setup and monitoring of the simulated climate change scenarios at TUMmesa. We are grateful to Sieglinde Sergl, Holger Paetsch, Luca Langlois, Paula Prucker, Phoebe Koppendorfer, Soizig Le Stradic, Veronika Kloska, and Yao Huang, who supported the experimental or data collection phase.

## Supporting information

**S1 File**. Description of study system

**S1 Table**. Species used in the mesocosm experiment to simulate urban grasslands

**S2 Table**. Composition of each experimental grassland community with the sown proportion of the two functional types ‘grasses’ and ‘forbs’

**S1 Fig**. Walk-in climate chambers of TUMmesa used for the mesocosm experiment with grasslands

**S2 Fig**. Realized daily air temperature, CO_2_ concentration, and relative humidity in the climate chambers

**S3 Fig**. Experimental design deployed in the climate chambers

**S3 Table**. Overview of indicator variables of urban grasslands measured in the mesocosm experiment

**S4 Fig**. Correlation matrix for the eight indicator variables of ecosystem functions assessed on mesocosm grasslands

**S5 Fig**. Clusters of seven indicator variables of ecosystem functions

**S4 Table**. Summary output of best models selected for describing drivers of single indicator variables of ecosystem functions of mesocosm grasslands

**S5 Table**. Multiple comparisons of forb proportion effects on single indicator variables of grassland functionality

**S6 Table**. Pairwise comparisons of forb proportion effects in interaction with climate-change related variables on plant hight of mesocosm grasslands

**S7 Table**. Output summary of best models for multifunctionality

**S2 File**. Multifunctionalty at 50% threshold of multifunctionality

## References

1. Gómez-Baggethun E, Gren Å, Barton DN, Langemeyer J, McPhearson T, O’Farrell P, et al. Urban ecosystem services. In: Elmqvist T, Fragkias M, Goodness J, Güneralp B, Marcotullio PJ, et al., editors. Urbanization, biodiversity and ecosystem services: challenges and opportunities: A global assessment. Dordrecht: Springer Netherlands; 2013. pp. 175–251.

2. IPCC, editor. Climate Change 2021: The physical science basis. Contribution of working group i to the sixth assessment report of the intergovernmental panel on climate change. Summary for policymakers. Cambridge University Press; 2021.

3. Gill S, Handley J, Ennos A, Pauleit S. Adapting cities for climate change: the role of the green infrastructure. Built Environment. 2007; 33:115–33. doi: 10.2148/benv.33.1.115.

4. Niemelä J, Saarela S-R, Söderman T, Kopperoinen L, Yli-Pelkonen V, Väre S, et al. Using the ecosystem services approach for better planning and conservation of urban green spaces: a Finland case study. Biodivers Conserv. 2010; 19:3225–43. doi: 10.1007/s10531-010-9888-8.

5. Norton BA, Coutts AM, Livesley SJ, Harris RJ, Hunter AM, Williams NS. Planning for cooler cities: A framework to prioritise green infrastructure to mitigate high temperatures in urban landscapes. Landscape and Urban Planning. 2015; 134:127–38. doi: 10.1016/j.landurbplan.2014.10.018.

6. Emilsson T, Ode Sang Å. Impacts of climate change on urban areas and nature-based solutions for adaptation. In: Kabisch N, Korn H, Stadler J, Bonn A, editors. Nature-based solutions to climate change adaptation in urban areas: linkages between science, policy and practice. Cham: Springer International Publishing; 2017. pp. 15–27.

7. Li D, Bou-Zeid E. Synergistic interactions between urban heat islands and heat waves: the impact in cities is larger than the sum of its parts. Journal of Applied Meteorology and Climatology. 2013; 52:2051–64. doi: 10.1175/JAMC-D-13-02.1.

8. Peñuelas J, Sardans J, Estiarte M, Ogaya R, Carnicer J, Coll M, et al. Evidence of current impact of climate change on life: a walk from genes to the biosphere. Glob Chang Biol. 2013; 19:2303–38. Epub 2013/03/25. doi: 10.1111/gcb.12143.

9. de Boeck HJ, Lemmens Cmhm, Vicca S, van den Berge J, van Dongen S, Janssens IA, et al. How do climate warming and species richness affect CO_2_ fluxes in experimental grasslands. New Phytol. 2007; 175:512–22. doi: 10.1111/j.1469-8137.2007.02122.x.

10. Fay PA, Hui D, Jackson RB, Collins HP, Reichmann LG, Aspinwall MJ, et al. Multiple constraints cause positive and negative feedbacks limiting grassland soil CO_2_ efflux under CO_2_ enrichment. Proc Natl Acad Sci U S A. 2021; 118. Epub 2020/12/21. doi: 10.1073/pnas.2008284117.

11. Polley HW, Jin VL, Fay PA. Feedback from plant species change amplifies CO_2_ enhancement of grassland productivity. Glob Chang Biol. 2012; 18:2813–23. Epub 2012/06/12. doi: 10.1093/aobpla/plv027.

12. Bloor JMG, Bardgett RD. Stability of above-ground and below-ground processes to extreme drought in model grassland ecosystems: Interactions with plant species diversity and soil nitrogen availability. Perspectives in Plant Ecology, Evolution and Systematics. 2012; 14:193–204. doi: 10.1016/j.ppees.2011.12.001.

13. Burri S, Niklaus PA, Grassow K, Buchmann N, Kahmen A. Effects of plant productivity and species richness on the drought response of soil respiration in temperate grasslands. PLoS ONE. 2018; 13:e0209031. Epub 2018/12/21. doi: 10.1371/journal.pone.0209031.

14. Onandia G, Schittko C, Ryo M, Bernard-Verdier M, Heger T, Joshi J, et al. Ecosystem functioning in urban grasslands: The role of biodiversity, plant invasions and urbanization. PLoS ONE. 2019; 14:e0225438. Epub 2019/11/22. doi: 10.1371/journal.pone.0225438.

15. Thompson GL, Kao-Kniffin J. Diversity enhances npp, n retention, and soil microbial diversity in experimental urban grassland assemblages. PLoS ONE. 2016; 11:e0155986. Epub 2016/05/31. doi: 10.1371/journal.pone.0155986.

16. Rudolph M, Velbert F, Schwenzfeier S, Kleinebecker T, Klaus VH. Patterns and potentials of plant species richness in high- and low-maintenance urban grasslands. Applied Vegetation Science. 2017; 20:18–27. doi: 10.1111/avsc.12267.

17. Decina SM, Hutyra LR, Gately CK, Getson JM, Reinmann AB, Short Gianotti AG, et al. Soil respiration contributes substantially to urban carbon fluxes in the greater Boston area. Environmental pollution. 2016; 212:433–9. Epub 2016/02/23. doi: 10.1016/j.envpol.2016.01.012.

18. Thompson GL, Kao-Kniffin J. Urban Grassland Management Implications for Soil C and N Dynamics: A Microbial Perspective. Frontiers in Ecology and Evolution. 2019; 7. doi: 10.3389/fevo.2019.00315.

19. Klaus VH, Kiehl K. A conceptual framework for urban ecological restoration and rehabilitation. Basic and Applied Ecology. 2021; 52:82–94. doi: 10.1016/j.baae.2021.02.010.

20. Norton BA, Bending GD, Clark R, Corstanje R, Dunnett N, Evans KL, et al. Urban meadows as an alternative to short mown grassland: effects of composition and height on biodiversity. Ecological Applications. 2019; 29:e01946. Epub 2019/07/22. doi: 10.1002/eap.1946.

21. Sehrt M, Bossdorf O, Freitag M, Bucharova A. Less is more! Rapid increase in plant species richness after reduced mowing in urban grasslands. Basic and Applied Ecology. 2019. doi: 10.1016/j.baae.2019.10.008.

22. Mata L, Andersen AN, Morán-Ordóñez A, Hahs AK, Backstrom A, Ives CD, et al. Indigenous plants promote insect biodiversity in urban greenspaces. Ecological Applications. 2021; 31:e02309. Epub 2021/04/07. doi: 10.1002/eap.2309.

23. Mody K, Lerch D, Müller A-K, Simons NK, Blüthgen N, Harnisch M. Flower power in the city: Replacing roadside shrubs by wildflower meadows increases insect numbers and reduces maintenance costs. PLoS ONE. 2020; 15:e0234327. Epub 2020/06/09. doi: 10.1371/journal.pone.0234327.

24. Manning P, van der Plas F, Soliveres S, Allan E, Maestre FT, Mace G, et al. Redefining ecosystem multifunctionality. Nat Ecol Evol. 2018; 2:427–36. Epub 2018/02/16. doi: 10.1038/s41559-017-0461-7.

25. Byrnes JEK, Gamfeldt L, Isbell F, Lefcheck JS, Griffin JN, Hector A, et al. Investigating the relationship between biodiversity and ecosystem multifunctionality: challenges and solutions. Methods in Ecology and Evolution. 2014; 5:111–24. doi: 10.1111/2041-210X.12143.

26. Meyer ST, Ptacnik R, Hillebrand H, Bessler H, Buchmann N, Ebeling A, et al. Biodiversity-multifunctionality relationships depend on identity and number of measured functions. Nat Ecol Evol. 2018; 2:44–9. Epub 2017/11/27. doi: 10.1038/s41559-017-0391-4.

27. Fry EL, Manning P, Allen DGP, Hurst A, Everwand G, Rimmler M, et al. Plant functional group composition modifies the effects of precipitation change on grassland ecosystem function. PLoS ONE. 2013; 8:e57027. Epub 2013/02/20. doi: 10.1371/journal.pone.0057027.

28. Mouillot D, Villéger S, Scherer-Lorenzen M, Mason NWH. Functional structure of biological communities predicts ecosystem multifunctionality. PLoS ONE. 2011; 6:e17476. Epub 2011/03/10. doi: 10.1371/journal.pone.0017476.

29. Fry EL, Savage J, Hall AL, Oakley S, Pritchard WJ, Ostle NJ, et al. Soil multifunctionality and drought resistance are determined by plant structural traits in restoring grassland. Ecology. 2018; 99:2260–71. Epub 2018/08/20. doi: 10.1002/ecy.2437.

30. van Sundert K, Arfin Khan MAS, Bharath S, Buckley YM, Caldeira MC, Donohue I, et al. Fertilized graminoids intensify negative drought effects on grassland productivity. Glob Chang Biol. 2021; 27:2441–57. Epub 2021/03/21. doi: 10.1111/gcb.15583.

31. Cantarel AAM, Bloor JMG, Soussana J-F. Four years of simulated climate change reduces above-ground productivity and alters functional diversity in a grassland ecosystem. Journal of Vegetation Science. 2013; 24:113–26. doi: 10.1111/j.1654-1103.2012.01452.x.

32. Bloor JMG, Pichon P, Falcimagne R, Leadley P, Soussana J-F. Effects of warming, summer drought, and co2 enrichment on aboveground biomass production, flowering phenology, and community structure in an upland grassland ecosystem. Ecosystems. 2010; 13:888–900. doi: 10.1007/s10021-010-9363-0.

33. Roy J, Picon-Cochard C, Augusti A, Benot M-L, Thiery L, Darsonville O, et al. Elevated CO_2_ maintains grassland net carbon uptake under a future heat and drought extreme. Proc Natl Acad Sci U S A. 2016; 113:6224–9. Epub 2016/05/16. doi: 10.1073/pnas.1524527113.

34. Giling DP, Beaumelle L, Phillips HRP, Cesarz S, Eisenhauer N, Ferlian O, et al. A niche for ecosystem multifunctionality in global change research. Glob Chang Biol. 2019; 25:763–74. Epub 2018/12/13. doi: 10.1111/gcb.14528.

35. Dietzel S, Rojas-Botero S, Fischer C, Kollmann J. Aufwertung urbaner Straßenränder als Anpassung an den Klimawandel und zur Förderung bestäubender Insekten. ANLiegen Natur. 2022; 44.

36. Deutscher Wetterdienst. Historical hourly station observations of 2m air temperature and humidity for Germany. Version v006. Retrieved from https://opendata.dwd.de. 2020.

37. Lan L, Chen J, Wu Y, Bai Y, Bi X, Li Y. Self-calibrated multiharmonic CO_2_ sensor using VCSEL for urban in situ measurement. IEEE Transactions on Instrumentation and Measurement. 2019; 68:1140–7. doi: 10.1109/TIM.2018.2863445.

38. Isbell F, Calcagno V, Hector A, Connolly J, Harpole WS, Reich PB, et al. High plant diversity is needed to maintain ecosystem services. Nature. 2011; 477:199–202. Epub 2011/08/10. doi: 10.1038/nature10282.

39. Lefcheck JS, Byrnes JEK, Isbell F, Gamfeldt L, Griffin JN, Eisenhauer N, et al. Biodiversity enhances ecosystem multifunctionality across trophic levels and habitats. Nat Commun. 2015; 6:6936. Epub 2015/04/24. doi: 10.1038/ncomms7936.

40. Maestre FT, Quero JL, Gotelli NJ, Escudero A, Ochoa V, Delgado-Baquerizo M, et al. Plant species richness and ecosystem multifunctionality in global drylands. Science. 2012; 335:214–8. doi: 10.1126/science.1215442.

41. Migliavacca M, Musavi T, Mahecha MD, Nelson JA, Knauer J, Baldocchi DD, et al. The three major axes of terrestrial ecosystem function. Nature. 2021; 598:468–72. Epub 2021/09/22. doi: 10.1038/s41586-021-03939-9.

42. Meeran K, Ingrisch J, Reinthaler D, Canarini A, Müller L, Pötsch EM, et al. Warming and elevated CO2 intensify drought and recovery responses of grassland carbon allocation to soil respiration. Glob Chang Biol. 2021; 27:3230–43. Epub 2021/05/06. doi: 10.1111/gcb.15628.

43. MacIvor JS, Lundholm J. Performance evaluation of native plants suited to extensive green roof conditions in a maritime climate. Ecological Engineering. 2011; 37:407–17. doi: 10.1016/j.ecoleng.2010.10.004.

44. MacIvor JS, Sookhan N, Arnillas CA, Bhatt A, Das S, Yasui S-LE, et al. Manipulating plant phylogenetic diversity for green roof ecosystem service delivery. Evol Appl. 2018; 11:2014–24. Epub 2018/09/28. doi: 10.1111/eva.12703.

45. Farrell C, Szota C, Williams NSG, Arndt SK. High water users can be drought tolerant: using physiological traits for green roof plant selection. Plant Soil. 2013; 372:177–93. doi: 10.1007/s11104-013-1725-x.

46. Dormann CF, Elith J, Bacher S, Buchmann C, Carl G, Carré G, et al. Collinearity: a review of methods to deal with it and a simulation study evaluating their performance. Ecography. 2013; 36:27–46. doi: 10.1111/j.1600-0587.2012.07348.x.

47. van der Plas F, Manning P, Allan E, Scherer-Lorenzen M, Verheyen K, Wirth C, et al. Jack-of-all-trades effects drive biodiversity-ecosystem multifunctionality relationships in European forests. Nat Commun. 2016; 7:11109. Epub 2016/03/24. doi: 10.1038/ncomms11109.

48. Zavaleta ES, Pasari JR, Hulvey KB, Tilman GD. Sustaining multiple ecosystem functions in grassland communities requires higher biodiversity. Proc Natl Acad Sci U S A. 2010; 107:1443–6. Epub 2010/01/04. doi: 10.1073/pnas.0906829107.

49. Barton K. MuMIn: Multi-Model Inference; 2020. Available from: https://CRAN.R-project.org/package=MuMIn.

50. Bates D, Mächler M, Bolker B, Walker S. Fitting linear mixed-effects models using lme4. Journal of Statistical Software. 2015; 67:1–48. doi: 10.18637/jss.v067.i01.

51. Brooks ME, Kristensen K, van Benthem KJ, Magnusson A, Berg CW, Nielsen A, et al. glmmTMB balances speed and flexibility among packages for zero-inflated generalized linear mixed modeling. The R Journal. 2017; 9:378–400. Available from: https://journal.r-project.org/archive/2017/RJ-2017-066/index.html.

52. Hartig F. DHARMa: Residual diagnostics for hierarchical (Multi-Level / Mixed) regression models; 2022. Available from: https://CRAN.R-project.org/package=DHARMa.

53. Lenth RV. emmeans: Estimated Marginal Means, aka Least-Squares Means; 2022. Available from: https://CRAN.R-project.org/package=emmeans.

54. R Core Team. R: A language and environment for statistical computing. Vienna, Austria; 2021. Available from: https://www.R-project.org/.

55. Song J, Wan S, Piao S, Knapp AK, Classen AT, Vicca S, et al. A meta-analysis of 1,119 manipulative experiments on terrestrial carbon-cycling responses to global change. Nat Ecol Evol. 2019; 3:1309–20. Epub 2019/08/19. doi: 10.1038/s41559-019-0958-3.

56. Zavalloni C, Vicca S, Büscher M, La Providencia IE de, Dupré de Boulois H, Declerck S, et al. Exposure to warming and CO_2_ enrichment promotes greater above-ground biomass, nitrogen, phosphorus and arbuscular mycorrhizal colonization in newly established grasslands. Plant Soil. 2012; 359:121–36. doi: 10.1007/s11104-012-1190-y.

57. Matthews HD. Implications of CO_2_ fertilization for future climate change in a coupled climate-carbon model. Glob Chang Biol. 2007; 13:1068–78. doi: 10.1111/j.1365-2486.2007.01343.x.

58. Pouyat RV, Szlavecz K, Yesilonis ID, Groffman PM, Schwarz K. Chemical, Physical, and Biological Characteristics of Urban Soils. In: Aitkenhead-Peterson J, Volder A, editors. Urban ecosystem ecology. Madison, WI, USA: American Society of Agronomy, Crop Science Society of America, Soil Science Society of America; 2010. pp. 119–52.

59. Armson D, Stringer P, Ennos AR. The effect of tree shade and grass on surface and globe temperatures in an urban area. Urban Forestry & Urban Greening. 2012; 11:245–55. doi: 10.1016/j.ufug.2012.05.002.

60. Armson D, Stringer P, Ennos AR. The effect of street trees and amenity grass on urban surface water runoff in Manchester, UK. Urban Forestry & Urban Greening. 2013; 12:282– 6. doi: 10.1016/j.ufug.2013.04.001.

61. Nagase A, Dunnett N. Amount of water runoff from different vegetation types on extensive green roofs: Effects of plant species, diversity and plant structure. Landscape and Urban Planning. 2012; 104:356–63. doi: 10.1016/j.landurbplan.2011.11.001.

62. Beard JB, Green RL. The role of turfgrasses in environmental protection and their benefits to humans. J Environ Qual. 1994; 23:452–60.

63. Lundholm J, MacIvor JS, Macdougall Z, Ranalli M. Plant species and functional group combinations affect green roof ecosystem functions. PLoS ONE. 2010; 5:e9677. doi: 10.1371/journal.pone.0009677.

64. Weisser WW, Roscher C, Meyer ST, Ebeling A, Luo G, Allan E, et al. Biodiversity effects on ecosystem functioning in a 15-year grassland experiment: Patterns, mechanisms, and open questions. Basic and Applied Ecology. 2017; 23:1–73. doi: 10.1016/j.baae.2017.06.002.

65. Marquard E, Weigelt A, Temperton VM, Roscher C, Schumacher J, Buchmann N, et al. Plant species richness and functional composition drive overyielding in a six-year grassland experiment. Ecology. 2009; 90:3290–302. doi: 10.1890/09-0069.1.

66. Boeck HJ de, Vicca S, Roy J, Nijs I, Milcu A, Kreyling J, et al. Global change experiments: Challenges and opportunities. BioScience. 2015; 65:922–31. doi: 10.1093/biosci/biv099.

67. Knapp AK, Smith MD, Hobbie SE, Collins SL, Fahey TJ, Hansen GJA, et al. Past, present, and future roles of long-term experiments in the LTER Network. BioScience. 2012; 62:377–89. doi: 10.1525/bio.2012.62.4.9.

